# RNA in extracellular vesicles during adolescence reveal immune, energetic and microbial imprints of early life adversity

**DOI:** 10.1101/2023.02.23.529808

**Authors:** L Korobkova, EL Morin, H Aoued, S Sannigrahi, KM Garza, ER Siebert, H Walum, RP Cabeen, MM Sanchez, BG Dias

**Author notes:** **Author Contributions**: ELM, BGD, and MMS conceived and planned the experiments. ELM and ERS collected and fractionated samples. HA, SS and KMG optimized the RNA extraction methods, and SS and ELM performed the RNA extraction. MMS designed the larger longitudinal developmental study that these animals were a part of and generated and characterized the animals. LK performed the analyses and interpreted results with ELM, HW, RPC and BGD. LK and BGD wrote the manuscript. All authors commented on the manuscript.

## Abstract

Exposure to early life adversity (ELA), including childhood maltreatment, is one of the most significant risk factors for the emergence of neuropsychiatric disorders in adolescence and adulthood. Despite this relationship being well established, the underlying mechanisms remain unclear. One way to achieve this understanding is to identify molecular pathways and processes that are perturbed as a consequence of childhood maltreatment. Ideally, these perturbations would be evident as changes in DNA, RNA or protein profiles in easily accessible biological samples collected in the shadow of childhood maltreatment. In this study, we isolated circulating extracellular vesicles (EVs) from plasma collected from adolescent rhesus macaques that had either experienced nurturing maternal care (CONT) or maternal maltreatment (MALT) in infancy. RNA sequencing of RNA in plasma EVs and gene enrichment analysis revealed that genes related to translation, ATP synthesis, mitochondrial function and immune response were downregulated in MALT samples, while genes involved in ion transport, metabolism and cell differentiation were upregulated. Interestingly, we found that a significant proportion of EV RNA aligned to the microbiome and that MALT altered the diversity of microbiome-associated RNA signatures found in EVs. Part of this altered diversity suggested differences in prevalence of bacterial species in CONT and MALT animals noted in the RNA signatures of the circulating EVs. Our findings provide evidence that immune function, cellular energetics and the microbiome may be important conduits via which infant maltreatment exerts effects on physiology and behavior in adolescence and adulthood. As a corollary, perturbations of RNA profiles related to immune function, cellular energetics and the microbiome may serve as biomarkers of responsiveness to ELA. Our results demonstrate that RNA profiles in EVs can serve as a powerful proxy to identify biological processes that might be perturbed by ELA and that may contribute to the etiology of neuropsychiatric disorders in the aftermath of ELA.

## 1. INTRODUCTION

The landmark collaboration between the CDC and Kaiser Permanente, and many follow-up studies have cemented our appreciation for the impact of Adverse Childhood Experiences (ACEs) on physical and mental health at later stages of life (Anda et al., 2010; Boullier & Blair, 2018; Chapman et al., 2004; Petruccelli et al., 2019). Among the many forms of ACEs or early life adversity (ELA), childhood maltreatment in the form of neglect and physical abuse is not only the most prevalent but is also a major risk factor for the emergence of adolescent and adult psychopathology (Brenhouse et al., 2018; Carr et al., 2013; Heim & Nemeroff, 2002; McLaughlin, 2020; Turecki et al., 2014; Vergara-Lopez et al., 2021). For example, compromised immune function, poor cardiovascular health and dysregulated emotional states seen in anxiety- and mood-related disorders, trauma- and stress-related disorders, as well as substance abuse disorders, manifest in the aftermath of childhood maltreatment (Cicchetti & Handley, 2019; Danese et al., 2007; Douglas et al., 2010; J. G. Johnson et al., 1999; Keyes et al., 2012; Lo & Cheng, 2007; McCrory et al., 2011; Nelson et al., 2017; Rodgers et al., 2004; Sinha, 2008; Teicher et al., 2003, 2016). Despite this well-established relationship between childhood maltreatment and the derailment of physiology and behavior, the mechanisms that contribute to such disruption are relatively unknown. Discovering these mechanisms is important for two reasons. First, to shed light on how childhood maltreatment might perturb biological systems to disrupt health in the future. Second, to identify biological signatures that could be used as biomarkers to determine who might be at risk of bearing the burden of maltreatment and who might escape the same. One way to achieve this objective is to profile changes in the (epi)genome, transcriptome and proteome in the aftermath of childhood maltreatment.

Most studies that have sought to determine biological signatures of ELA have profiled changes in DNA methylation in blood cells (Catale et al., 2020; Khulan et al., 2014; Kinnally et al., 2011; Nieratschker et al., 2014; Romens et al., 2014; Szyf, 2013). From these powerful approaches, we have come to appreciate that different kinds of ELA result in unique signatures of DNA methylation in blood cells and that molecular signatures in peripheral systems like the hematopoietic system do in fact change in response to ELA. However, despite this progress, because ELA encapsulates many forms of stress and the hematopoietic system is but one biological system in the body, we have limited appreciation for how the entire organism is specifically impacted by bouts of maltreatment during infancy. Needed to fill these gaps in our knowledge are a biological sample that could be a proxy for biological perturbations that span the entire organism and is not focused on specific cells, tissues, organs and systems. Additionally, an unfortunate but ideal experimental framework within which to ask these questions would be one in which childhood maltreatment is the form of ELA that infants experience, and in which the maltreatment encapsulates both, neglect and physical abuse, without other confounding factors like prenatal exposure to drugs, lack of access to proper nutrition and medical care. To satisfy these criteria and gain better insights into the etiology and biomarkers of maltreatment-exacerbated disruptions to physiology and behavior, we used a highly translational and naturally occurring model of infant maltreatment in which rhesus macaques are recipients of both, physical abuse and neglect (in the form of infant rejection), from mothers during infancy (Drury et al., 2017; K. M. McCormack et al., 2022). Next, during adolescence we profiled RNA found in extracellular vesicles (EVs) circulating in the plasma of animals that had experienced infant maltreatment and compared this profile to RNA in EVs extracted from animals reared with nurturing maternal care.

EVs are nanometer-scale membrane vesicles secreted by most cell types and can be found in biofluids (Alberro et al., 2021; Arraud et al., 2014; Brenna et al., 2021; Caby et al., 2005; Holm et al., 2018; Mustapic et al., 2017; van der Pol et al., 2012; Yáñez-Mó et al., 2015; Yates et al., 2022). Important agents of inter-cellular communication and interaction (Abey et al., 2017; Bischoff et al., 2022; Casado-Díaz et al., 2020; Delpech et al., 2019; Desdín-Micó & Mittelbrunn, 2017; Diaz-Garrido et al., 2021; Gabisonia et al., 2022; Moreno-Gonzalo et al., 2018; Ramachandran & Palanisamy, 2012; Saeedi et al., 2019; Vojtech et al., 2014; Wahlund et al., 2017), EV cargo is affected by ELA in humans and rodents (Allen & Dwivedi, 2020; Alshanbayeva et al., 2021; Morrison et al., 2022; Pinson et al., 2021; Ran et al., 2022). Based on this background, we hypothesized that the biological content of EVs would serve as powerful indicators of how organism-wide biology gets coordinated at the molecular level to respond to the affront of infant maltreatment. With EVs containing not only fragments of peptides and signal molecules, but also RNAs, the identity and abundance of these RNAs could serve as a valuable window into how maltreatment impacts biology. With this motivation, we sought to profile total RNA found in circulating EVs in plasma of adolescent macaques that had either experienced maternal maltreatment in infancy (MALT) or nurturing maternal care (CONT).

## 2. METHODS

### 2.1 Subjects

Thirteen adolescent (seven males and six females) rhesus macaques (*Macaca mulatta*) were included in this study. These macaques were born and raised with their mothers and families in complex social groups at the Emory National Primate Research Center (ENPRC) Field Station. Social groups consisted of 75-150 adult females, their sub-adult and juvenile offspring, and 2-3 adult males. Animals were housed in outdoor compounds, with access to climate-controlled indoor areas. Standard high fiber, low fat monkey chow (Purina Mills Int., Lab Diets, St. Louis, MO) and seasonal fruits and vegetables were provided twice daily, in addition to enrichment items. Water was available ad libitum.

Part of a larger longitudinal study of the effects of ELA, the animals in our study were well-characterized throughout infancy and the pre-pubertal period (Drury et al., 2017; Howell et al., 2013, 2017; K. M. McCormack et al., 2022; Morin et al., 2020). Six of the subjects (3 males and 3 females) experienced maternal maltreatment (MALT), and the other seven (4 males and 3 females) received nurturing maternal care (CONT). Infant maltreatment is defined by co-morbid experience of maternal physical abuse, rejection and neglect of the infant during the first three months of life, resulting in pain, emotional distress, and elevations in stress hormones (Drury et al., 2017; Howell et al., 2013, 2017; Maestripieri et al., 2000; Maestripieri & Carroll, 1998; McCormack et al., 2006, 2009, 2015, 2022). Animals in the CONT group were the recipients of positive caregiving behaviors like cradling and grooming and none of the physical abuse or rejection experienced by MALT animals. Each infant in this study was randomly assigned at birth to be cross-fostered by a CONT or MALT foster mother. This was done to disentangle and control for effects of prenatal and heritable factors that may confound the effects of ELA in the larger developmental study, where groups were also counterbalanced by social dominance status and infants were assigned from different matrilines and paternities to provide genetic and social diversity (Drury et al., 2017; Howell et al., 2017; Morin et al., 2020).

All 13 animals included in this study were cross-fostered to a foster mother of the same biological group (CONT or MALT); therefore, the prenatal and postnatal environments were congruent. More specifically, an infant born to a mother that was known to demonstrate MALT was raised by another female who exhibited MALT to her offspring and an infant born to a mother that provided CONT care to her infants was fostered by a female who also provided CONT care. Behavioral measures of maternal care were collected during the first 3 postnatal months to characterize competent care received by CONT infants, and to quantify rates of maternal physical abuse and rejection rates received by MALT infants. A more detailed description of the infant rhesus maltreatment model and behavioral methods can be found in (Drury et al., 2017; Howell et al., 2017; K. M. McCormack et al., 2022). Interestingly, similar rates of abuse and rejection are exhibited by maltreating foster mothers towards foster infants than towards previous biological infants. Control mothers did not exhibit physical abuse or rejection towards cross-fostered infants from biological Control or Malt mothers in the larger developmental study. This suggests that maltreatment is a maternal trait, and not triggered by the infant.

At approximately 4 years of age, these animals were transferred to the ENPRC Main Station. Upon arrival, animals were pair-housed in indoor home cages and fed Purina monkey chow (Ralston Purina, St. Louis, MO, USA), supplemented with fruit and vegetables daily, and water was available ad libitum. Environmental enrichment was provided on a regular basis. The colony is maintained at an ambient temperature of 22 ± 2°C at 25-50% humidity, and the lights set to a 12-h light/dark cycle (lights on at 7h; lights off at 19h). The animals were given several months to acclimate to the move and new housing environment before subsequent experimental phases began.

All experimental procedures and animal care were in accordance with the Animal Welfare Act and the U.S. Department of Health and Human Services “Guide for the Care and Use of Laboratory Animals” and approved by the Emory Institutional Animal Care and Use Committee (IACUC).

### 2.2 Plasma Collection

Blood samples were collected mid-adolescence (4.61± 0.42 years old) while the animals were sedated (telazol 5mg/kg, i.m.) in preparation for MRI or PET imaging. Blood was drawn from the femoral vein into a vacutainer CPT tube, kept on ice, and cold centrifuged as soon as possible. Plasma was pipetted from the top of the fractionated sample and aliquots were stored at -80°C.

### 2.3 Isolation of EVs

Plasma samples were first thawed and thrombin was added to the plasma to a final concentration of 5U/mL to defibrinate the plasma. After centrifugation, EVs were isolated from the resulting supernatant using the ExoQuick Exosome Isolation kit from System Biosciences following manufacturer’s instructions. Purity of EV collection was ascertained by Western Blotting for EV-marker CD-9 **(Fig. S1A)**.

### 2.4 RNA extraction & sequencing

RNA was extracted from EVs using a standard guanidinium thiocyanate-phenol-chloroform, or TRIzol® (Thermo Fischer Scientific, Waltham, MA) extraction protocol, which has been shown to be a robust method for total RNA extraction (Mraz et al., 2009) **(Fig. S1B)**. Extracted RNA was stored at -80°C prior to sequencing. Total RNA sequencing, including stranded library prep and treatment with Ribogone™ (Takara Bio USA Inc., Mountain View, CA) was performed by the Emory National Primate Research Center Genomics Core on the Illumina HiSeq 3000 system (single-end strand specific reads, 100 bp).

### 2.5 Bioinformatics analysis

Reads were trimmed, aligned and counted with STAR algorithm (Dobin et al., 2013) using the rhesus macaque Mmul10 genome annotation (INSDC Assembly GCA_003339765.3) published by The Genome Institute at Washington University School of Medicine in 2019. Summary mapping statistics were calculated with STAR for each sample **(Fig. S2)**. Coverage was computed using the bedtools coverage function on each stranded .bam file. To ensure robustness, like we did in Morin et al., 2022, our analyses were performed on genes on the sense strand with >= 80% coverage.

### 2.6 Differential expression analysis

To identify differentially expressed genes in control and maltreated groups, we used Limma-Voom R package (R Studio version 1.3.959) (Law et al., 2014) to examine group differences and a model which accounted for sex as a covariate **(Table 1, Fig. S3)**.

**Table 1:** Infant maltreatment alters RNA found in EVs. Differential gene expression (DGE) analysis was performed to identify RNA that is differentially expressed in EVs from CONT and MALT animals. After correcting for multiple comparisons, RNA associated with 11 genes were found to be differentially expressed in EVs from MALT animals compared to CONT samples. RPL23A, RPL26, RPS27A - Ribosomal Protein L23a WSCD1 - sulfotransferase activity IDH1 - Peroxisomes component, isocitrate dehydrogenase 1. ACTB - beta (β)-actin CD74 - MHC II chaperone, antigen presentation GMFG - Glia Maturation Factor Gamma GALM - Galactose Mutarotase, maintenance of the equilibrium between anomers of galactose LAGE3 - transfer of the threonylcarbamoyl moiety of threonylcarbamoyl-AMP (TC-AMP) ATP13A5 - nucleotide binding and ATPase-coupled cation transmembrane transporter activity

| geneID | logFC | AveExpr | t | adj.P.Val |
| --- | --- | --- | --- | --- |
| *rpl23a* | -4.629 | 9.993 | -5.268 | 0.046 |
| *wscd1* | -7.705 | 1.161 | -5.136 | 0.046 |
| *idh1* | 7.401 | 2.981 | 5.105 | 0.046 |
| *rpl26* | -5.563 | 7.517 | -5.033 | 0.046 |
| *actb* | -3.587 | 10.420 | -4.985 | 0.046 |
| *cd74* | -4.722 | 7.693 | -4.834 | 0.046 |
| *gmfg* | -8.006 | 4.420 | -4.834 | 0.046 |
| *rps27a* | -3.190 | 9.669 | -4.804 | 0.046 |
| *galm* | 7.343 | 3.160 | 4.760 | 0.046 |
| *lage3* | -7.832 | -0.028 | -4.667 | 0.047 |
| *atp13a5* | 7.657 | 2.725 | 4.623 | 0.047 |

### 2.7 GO analysis

Gene Ontology (GO) Enrichment Analyses (Package clusterProfiler version 4.0.2, GO.db_3.13.0) were performed with biological processes gene sets within the *Macaca mulatta* reference list to identify broad cellular and molecular pathways where RNA in EVs may play a role. Functional enrichment analysis was performed with R package clusterProfiler on features with an absolute logFC >2 and p<0.05 **(Fig. 1, Fig. S4)**.

**Fig. 1:**
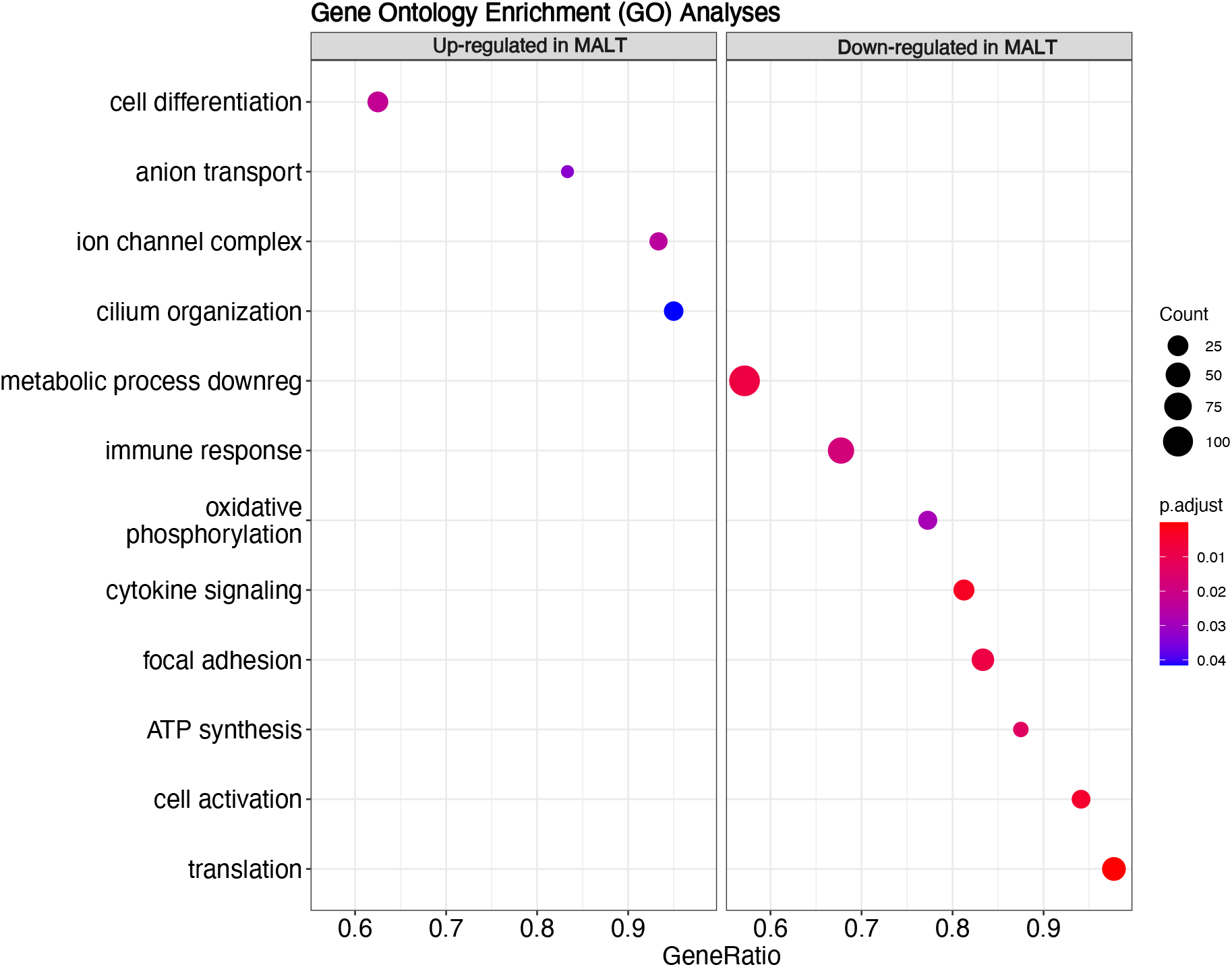
Perturbations of biological processes by infant maltreatment are evident from RNA signatures found in circulating EVs. Gene Ontology (GO) analysis of differential RNA expression between the RNA cargo of CONT and MALT EVs revealed upregulated (left panel) and downregulated (right panel) RNA in EVs of the MALT group relative to the CONT group. Data demonstrate that RNA cargo associated with immune, cellular energetics, and translation are the most differentially regulated in MALT EVs. Each row represents a different term from the enriched Gene Ontology for BP (biological process). The position of each dot represents the gene ratio (the percentage of total Differentially Expressed Genes in the GO term). The color of each dot represents adjusted p-value (with red being the smallest), and the size reflects the number of genes belonging to a given group. Final list of significant GO terms was manually curated to remove functional duplicates which share majority of genes. Full list is provided in **Table S1**.

### 2.8 Metatranscriptomics to analyze microbiome-related transcripts

Given the large percentage of unmapped reads to the macaque genome, we performed metatranscriptomic analysis on those unmapped reads, using the Kraken algorithm to map genomic reads to the bacterial and viral genome (Lu et al., 2022; Wood & Salzberg, 2014). To test robustness of this approach we also performed the classification on raw data. We observed significant increase in the percentage of reads identified as microbiome-related when we excluded reads that aligned to reference Macaque genome **(Fig. S5)**. The Bracken algorithm (Lu et al., 2017) was then used to determine the relative abundance of bacterial species in CONT and MALT groups. To test whether the depth of the sequencing was enough to correctly reconstruct the microbial signatures, we randomly sub-sampled reads from Bracken output and plotted the rarefaction curve **(Fig. S6)**. Species were retained for the class- and phyla-level composition analysis. To determine which bacteria were common in control and maltreated groups, we curated the final species’ list to include only those species that were present in all samples.

To measure the discordance between microbiome profiles in EVs from CONT and MALT animals, we developed a statistical approach using permutation testing. As a test statistic, we chose a measure of the total number of species that were “different” between groups. We defined “different” as having a species in at least 4 samples in one group, but not the other, because 4 is the majority in the smaller CONT group (n in CONT group = 6). Specifically, for each species we counted how many CONT and MALT cases had that species, and then determined a difference by absolute (n of presence in CONT - n of presence in MALT) > 4. We then counted how many bacterial species were different based on this criteria. Our permutation test built a null distribution of the total number of different species across 5000 iterations, and we computed a single p-value as the probability of getting a value as extreme (large) as our observed count under this null distribution **(Fig. S7)**.

To identify class signatures that are common for both groups and predominant in one, we grouped species with taxize R package. **(Fig. 2)**. The difference between CONT and MALT groups in microbiome species number and RNA abundance of the microbiome community was analyzed with Wilcoxon rank sum test **(Fig. 3, Fig. S8)**.

**Figure 2:**
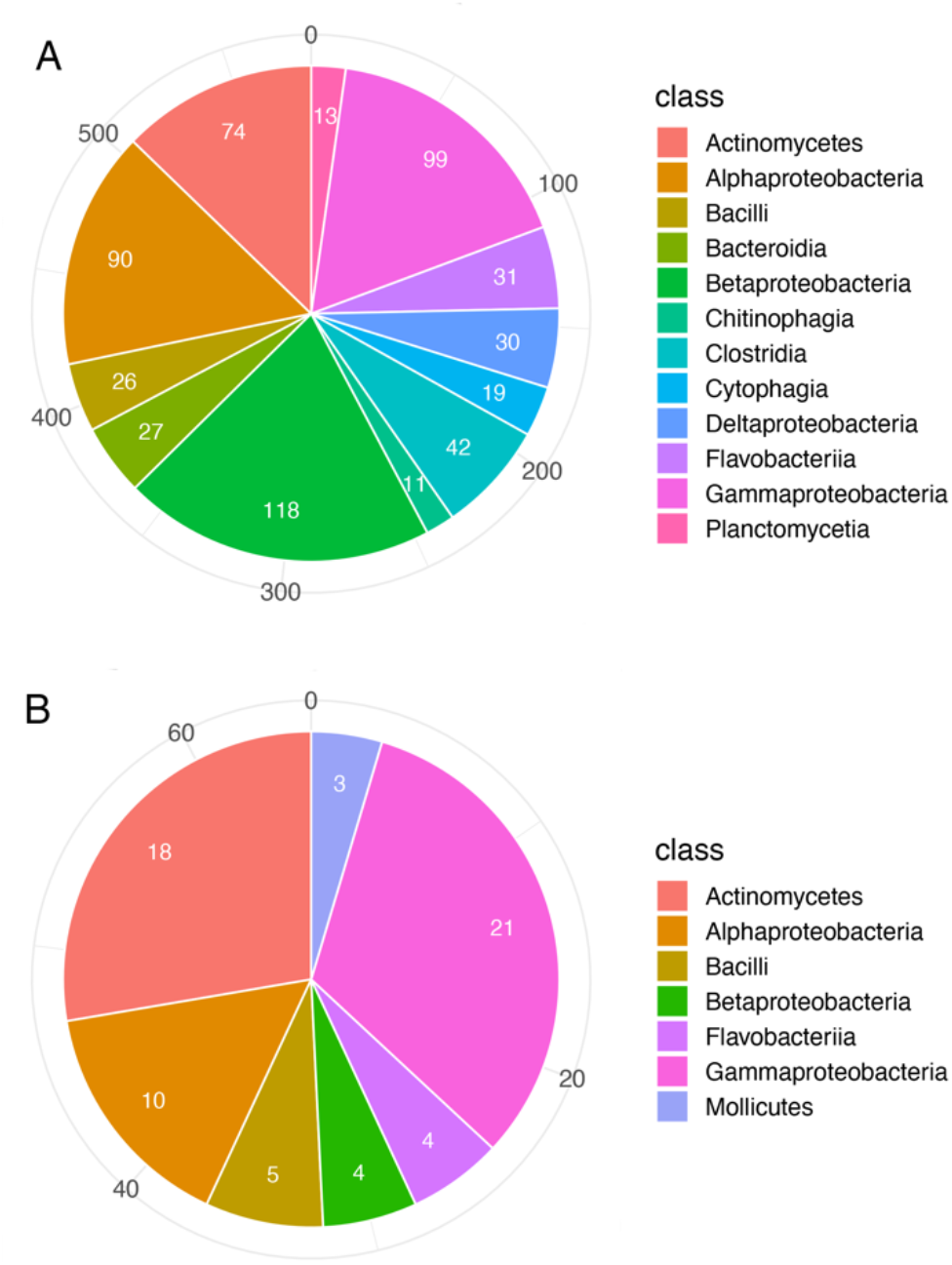
Microbiome signatures in EVs from CONT and MALT samples as detected by RNA content in circulating EVs. **(A)** Pie chart showing classes of microorganisms that have RNA signatures in EVs of both CONT and MALT groups based on alignments of RNA found in EVs to the meta-transcriptome (only classes that have species > 10 are shown). **(B)** Pie chart showing classes of microorganisms that are more prevalent in EVs of the CONT group as detected by RNA signatures in EVs (five classes with n<3 species are not shown). Numbers inside wedges show how many species were detected in each class. Numbers on the circumference reflect cumulative count of the species across all classes.

**Figure 3:**
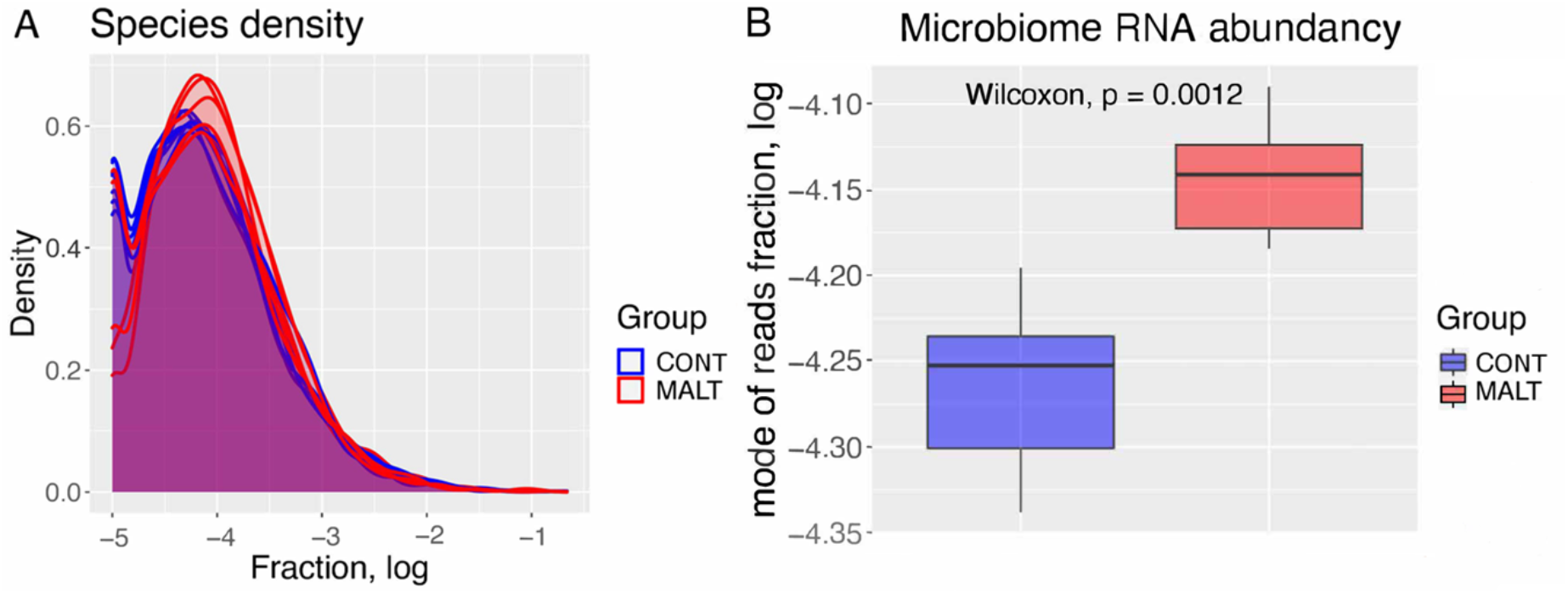
RNA in EVs reflect changes in microbiome diversity after infant maltreatment. **(A)** Microbial species density in CONT and MALT groups. Density is plotted as a function of the fraction of RNA reads on a log scale. Each line represents one sample and is color-coded by CONT or MALT group. Qualitatively, the plot shows the MALT samples are skewed to the right and have relatively greater peak values demonstrating greater RNA abundance associated with the microbiome in the EVs of MALT animals. **(B)** Plot showing the difference in modes (peak values) of microbial RNA abundancy in CONT (blue) and MALT (red) groups. The boxplots represent the median (line in the middle of the box), upper 25% quantile, lower 25% quantile, and the interquartile ranges (upper and lower whiskers). Wilcoxon test derived p-value is shown. On average, peak values of RNA abundancy in MALT group are greater than in CONT (modes are shown on a log scale) indicating that RNA signatures associated with the microbiome are altered by infant maltreatment.

### 2.9 Stool 16S rRNA analysis workflow

We sought to compare microbiome composition in stool and EVs in order to understand the possible source of microbiome-associated RNA in EVs. Stool samples were collected opportunistically and stored at -80 °C prior to DNA extraction with the E.Z.N.A. Stool DNA Kit (Omega Bio-tek, Inc), including a second incubation to isolate gram-positive bacteria. The samples were sequenced with the Illumina MiSeq system. This sequencing protocol used primer pair sequences for the V4 region of the 16S rRNA gene, creating amplicons that are approximately 460 base pairs (bp) in length. Bioconductor workflow in R was used to trim, filter, and test this data using the packages dada2 and phyloseq.

### 2.10 Comparison between microbiome signatures found in stool samples and EVs

Microbiome species noted in both, EVs and stool, were classified according to their phyla using the methods described in Sections 2.8 and 2.9. First for each specimen and phylum, we computed fractional values by dividing number of detected RNAs and 16S rRNA gene copies in EVs or stool, by the sum total of their copies in that sample. Next, data were averaged by group and specimen type. Abundance values were transformed to relative abundance, and the top five most abundant phyla in every group were included in the final comparison and plotted (**Fig. 4, Table 2**).

**Figure 4:**
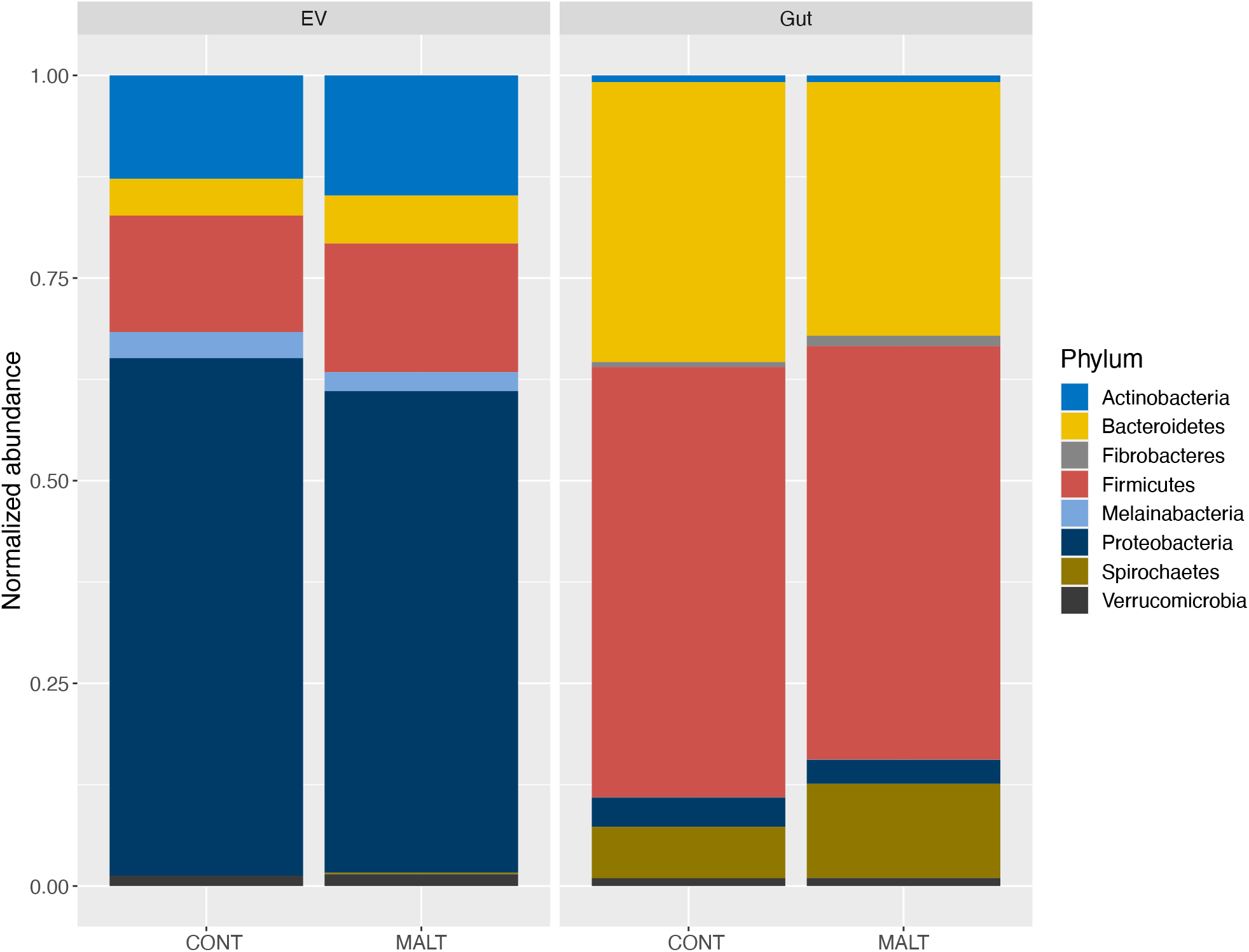
Comparison of microbiome-associated molecular signatures in plasma-derived circulating EVs and the stool-derived gut microbiome suggest different sources of origin. Plot showing the fraction of most abundant phyla in plasma-derived EVs and gut microbiome calculated from RNA and 16S rRNA sequencing, respectively, in CONT and MALT groups. Data are shown as average across samples within groups. Each phylum is color-coded. Firmicutes and Bacteroidetes are the most abundant phyla in gut microbiome. Proteobacteria are the most abundant phyla detected in plasma-derived EVs.

**Table 2:** Overlap in most abundant phyla found in plasma-derived EVs and gut-derived samples with different proportions. Table showing abundance of eight major phyla averaged across samples and groups. Bacteroidetes, Firmicutes and Proteobacteria are three most abundant taxa across all four groups of comparison.

| Phylum | Normalized abundance, % |  |  |  |
| --- | --- | --- | --- | --- |
|  | EV |  | Gut |  |
|  | CONT | MALT | CONT | MALT |
| Actinobacteria | 0.1276 | 0.1483 | 0.0084 | 0.0084 |
| Bacteroidetes | 0.0453 | 0.0589 | 0.3452 | 0.3128 |
| Fibrobacteres | - | - | 0.0062 | 0.0128 |
| Firmicutes | 0.1436 | 0.1589 | 0.5309 | 0.5102 |
| Melainabacteria | 0.0322 | 0.0234 | 0.0002 | 0.0003 |
| Proteobacteria | 0.6385 | 0.5941 | 0.0361 | 0.0296 |
| Spirochaetes | - | 0.0018 | 0.0633 | 0.1161 |
| Verrucomicrobia | 0.0127 | 0.0146 | 0.0096 | 0.0099 |

### 2.11 Data availability

Sequencing data has been deposited on the GEO database and can be accessed via GEO Accession #: GSE223984 (EV) and GSE225873 (stool microbiome). Analysis code is available at https://github.com/BDiasLab/NHP_exo.

## 3. RESULTS

### 3.1 Differential Expression Analysis of RNA content in plasma-derived EVs

Differential expression analysis of reads located on the sense strand revealed genes that were both, increased and suppressed in MALT compared to CONT animals. 11 genes survived FDR multiple comparison correction with adjusted p-value < 0.05 (**Table 1, Fig. S3)**.

### 3.2 GO analysis of RNA content in plasma-derived EVs

GO Enrichment Analysis on 908 genes with an absolute logFC >2 and p-value <0.05 revealed functional groups of genes that were significantly activated (upregulated) and suppressed (downregulated) in the MALT group compared to CONT. These differentially altered functional sets of genes are related to translational processes, ATP energetics, mitochondrial organization, immune function, epithelial cell differentiation, metabolic processes and oxidative phosphorylation **(Fig. 1, Fig. S4, Table S1)**.

### 3.3 RNA in plasma-derived EVs align to microbial transcriptome

Summary mapping statistics revealed that 29.9% to 76.83% of reads (unmapped and mapped to multiple loci) did not map to reference Mul10 genome **(Fig. S2)**. The percentage of genes with coverage (>=80%) that did align to Mul10 genome was on average 65.52%. To explore the identity of the reads that remained unmapped to the Mul10 genome, we hypothesized that they might belong to the microbial metatranscriptome. Running Kraken analysis using reads that did not align to the Mmul10 genome resulted in a high percentage of reads aligned to MiniKraken database, confirming that the microbial transcriptome makes up a substantial proportion of the RNA content in EVs **(Fig. S5)**. Rarefaction curve (Gart et al., 1982) revealed that our sequencing depth was sufficient to detect the rarest species **(Fig. S6)**.

### 3.4 Prevalence of RNA signatures associated with the microbiome in plasma-derived EVs of CONT and MALT animals

RNA found in EVs from CONT and MALT animals revealed 691 species to be present in both these groups **(Fig. 2**, species list included in **Table S2)**. Further examination of bacterial and viral classes revealed that Actinomycetia, Alphaproteobacteria, Betaproteobacteria and Gammaproteobacteria were the predominant clades in both groups **(Fig. 2A)**. Permutation revealed that 81 species were differentially prevalent across the groups **(Fig. S7)**. Specifically, species belonging to Actinomycetia, Alphaproteobacteria and Gammaproteobacteria were found to be more prevalent in CONT animals than in MALT animals **(Fig. 2B)**. Plasma EVs from MALT animals had RNA signatures of *Bacillus sp. X1* and *Rhodococcus opacus* that were not detected in CONT animals. Plasma EVs from CONT animals in turn had RNA signatures of several species (*Anabaena sp. 90, Bordtella hinzii, Bradyrhizobium diazoefficiens, Burkholderia sp. CCGE1003, Candidatus nitrosomarinus Catalina, Klebsiella quasipneumoniae, Krypidia sporamanii, Nonlabens marinus, Pseudomonas orientalis, Pseudomonas palleroniana, Pseudomonas parafulva, Rickettsia amblyommatis, Staphylococcus xylosus, Streptomyces globisporus, Streptomyces leeuwenhoekii, Tsukamurella paurometabloa*) that were not detected in MALT animals.

### 3.5 Microbiome richness and RNA abundance in plasma-derived EVs

RNA profiling of microbiome signatures in circulating EVs did not reveal any significant differences in the total number of microbiome species between CONT and MALT samples. **(Fig. S8)**. Analysis of the distribution of the RNA fraction profile revealed different patterns of microbiome RNA abundance between the CONT and MALT groups (**Fig. 3A)**. The MALT group showed on average higher mode of fraction of RNA reads in comparison to CONT samples (p-value = 0.0012, Cohen’s d effect size = 2.78) **(Fig. 3B**) demonstrating that the MALT group had a distribution of RNA abundancy in EVs associated with the microbiome that was distinct from CONT group.

### 3.6 Determining origins of the RNA in plasma-derived EVs

To compare microbiome-associated molecular signatures in EVs isolated from circulating plasma and the microbiome of the gut (using stool as a proxy), we chose to compare the five most abundant phyla in each sample (Plasma EV CONT, Plasma EV MALT, Gut CONT and Gut MALT). While this selection criteria resulted in Bacteroidetes, Firmicutes and Proteobacteria being common for all four groups, their proportions in Plasma EV-vs Gut-derived samples **(Fig. 4 and Table 2)** were different. Specifically, Firmicutes is the most abundant phylum in the gut (on average 50%), while Proteobacteria is most abundant in Plasma EVs (on average 60%). Bacteroidetes, the second most abundant phylum in gut-derived samples (on average 33%), makes up only 5% in plasma-derived EVs. Spirochaetes that is more abundant in MALT than in CONT in gut microbiota comprises a small percentage of the microbiota profile in plasma-derived EVs in MALT animals and cannot be detected in CONT samples after performing a median split on raw data.

## 4. DISCUSSION

EVs present an exciting frontier in our understanding of how stress comes to be embedded in our biology (Beninson & Fleshner, 2014; F. Chen et al., 2021; de Jong et al., 2012; Fleshner & Crane, 2017; Lin et al., 2015; Luarte et al., 2017; Smythies et al., 2014; Sung et al., 2021). By virtue of being released from cells that make up all organs and tissues (Kalluri & LeBleu, 2020; Kowal et al., 2014; Li et al., 2020; Pegtel & Gould, 2019; Properzi et al., 2013; Simons & Raposo, 2009; Théry et al., 2002; van Niel et al., 2006), their contents could serve as a harbinger of the processes and pathways that come to be disrupted from exposure to ELA. A corollary to this perspective is that their contents could also provide clues about processes that might result in the manifestation of physiological and behavioral disruptions in the aftermath of ELA. With this intent, we used a translational NHP model of ELA resulting from naturally occurring infant maltreatment (Howell et al., 2013, 2017; K. McCormack et al., 2006, 2009; K. M. McCormack et al., 2022; Morin et al., 2019, 2020, 2022; Wakeford et al., 2018) and profiled total RNA in circulating plasma EVs obtained from adolescent macaques that had received nurturing care (CONT) or maltreatment (MALT) from their mothers as infants. From these EV-associated RNA signatures, our results suggest that perturbations of the immune system, cellular energetics and translation ought to be considered as biomarkers of infant maltreatment and possible mediators of detrimental physiological and behavioral consequences of such maltreatment. Of equal importance is our finding that maltreatment alters microbiome species diversity and RNA abundance as assayed by microbiome-associated RNA in circulating EVs.

Broadly, our GO enrichment analysis results are in congruence with studies that have probed immune system-related and energetic consequences of stress. Immune function and cellular energetics via mitochondrial function appear to be important processes in this context because they are perturbed across species after exposure to childhood maltreatment. Social subjugation and consequent low social status in wild baboon populations impacts proportion of immune cells and altered gene expression in helper T cells, cytotoxic T cells, B cells, monocytes and Natural Killer cells (Snyder-Mackler et al., 2016). Maltreatment rates experienced in infancy were associated with increased activation in pro-inflammatory pathways in monocytes of juvenile macaques (Sanchez et al., 2007). Chronic social defeat in rodents has been shown to trigger inflammation and impact immune function (Niraula & Sheridan, 2019; Powell et al., 2013). Compromised immune function and associated inflammation have also been discussed as strong mediators of the pathological influences of childhood maltreatment (Baldwin et al., 2018; Coelho et al., 2014; Cohen-Woods et al., 2018; Danese et al., 2007; do Prado et al., 2017; Gonzalez, 2013; Osborn & Widom, 2020; O’Shields et al., 2022). A more stringent analysis of our data using DGE also pointed to immune function as a biological process that was perturbed after maltreatment. RNA aligning to *cd74* that is expressed by B lymphocytes and antigen presenting cells and induces a signaling cascade that results in regulation of cell proliferation and survival, as well as RNA aligning to *gmfg* (Glia Maturation Factor Gamma - a cytokine-responsive protein) were found to be present in lower levels in EVs from MALT animals.

Cellular energetics driven by mitochondrial structure and function are also becoming an increasingly appreciated contributor to the pathophysiological consequences of early life stress (Picard et al., 2018; Zitkovsky et al., 2021). ELA in humans has been shown to alter mitochondrial DNA copy number (Tyrka et al., 2016) and ATP-related mitochondrial bioenergetics (Gumpp et al., 2022). Exposure to the highly influential limited nesting and bedding model of early life stress in mice (Gilles et al., 1996; Molet et al., 2014) altered mitochondrial function in the brain and liver (Eagleson et al., 2020). Similar results on mitochondrial biology were obtained in juvenile + adult double-hit model of stress in rats (Nold et al., 2019). DGE identified RNA aligning to genes associated with energetic process were upregulated in MALT EVs. Specifically, we found that RNA aligning to *idh1* (breaks down fats for energy and protect cells from harmful molecules; mutations are associated with gliomas; mutant IDH1 confers resistance to energy stress through PFKP-induced aerobic glycolysis and AMPK activation), *galm* (converts galactose to glucose), and to *atp13a5* (cation-transporting ATPase) were upregulated in MALT EVs. The increased representation of these RNA in EVs might suggest a compensatory effect to balance out the dysregulation in mitochondrial function that may occur after exposure to maltreatment and that plays the major role in energetic balance of every cell.

Our data demonstrate that RNA found in EVs could be as reflective of perturbations to these systems after maltreatment as are disruptions of the direct actors and actions of immune function like the proportion of immune cells and signatures of inflammation like cytokines and of cellular energetics by measurement of metabolites and mitochondrial function. Our work demonstrating imprints of maltreatment on immune and cellular energetics sets the stage from which we ought to consider probing interactions between these mitochondrial function in immune cells as a means by which maltreatment leaves multi-dimensional biological scars in offspring. Doing so would be in keeping with the attention that mitochondrial dynamics in immune cells are getting as one way via which alterations in cellular energetics might impact immune function (Faas & de Vos, 2020; Mehta et al., 2017; Rambold & Pearce, 2018; Xie et al., 2020).

Protein translation is a critical component of cellular function. Disruptions of gene expression associated with the same have been observed in human biological samples probed in the context of neurodegeneration (Johnson et al., 2020; Seyfried et al., 2017; Wingo et al., 2021). Less is known about the potential for alterations in ribosomal assembly, function and translation after a history of childhood maltreatment. Our GO analyses highlighted that RNA associated with translational processes and DGE data identified *rpl23a, rpl26, rps27a* ribosomal-associated RNA to be less present in our MALT samples. Taken together, these complementary approaches suggest that translation might be a process and pathway that ought to be studied as a biomarker of ELA and a potential mediator of its psychopathology. The only evidence of this relationship that we could find in the context of early life adversity was the report that exposure to brief daily separation, a model of ELA in mice, resulted in decreased availability of ribosomal RNA in the hippocampus and this was posited to blunt hippocampal growth after this form of ELA (Wei et al., 2014).

The relatively low percentage of our sequencing results uniquely aligning with the macaque genome was puzzling. However, given that EVs bleb off from all cells in an organism, we posited that the RNA in EVs might originate from sources independent of the mammalian host. With bacterial and viral communities occupying major niches in the biological systems of hosts, we thought to align our sequencing results with the microbial meta-transcriptome using the Kraken algorithm. While Kraken is typically used to map DNA sequences to the bacterial genome, studies suggest the utility of using it to align RNA sequences to the same (Chen et al., 2018; Flygare et al., 2016; Neves et al., 2017; Purcell et al., 2017; Reck et al., 2015; Swanson et al., 2020; Tarallo et al., 2019). From these analyses, we were surprised to find that the reads that did not uniquely map to the macaque genome showed high alignment with the meta-transcriptome of prokaryotic microorganisms. This finding is congruent with studies showing that microbiome-associated RNA has been found in circulating exosomes in humans (Cuesta et al., 2021; Diaz-Garrido et al., 2021; Jones et al., 2021; ñahui Palomino et al., 2021; Schaack et al., 2022; Tsatsaronis et al., 2018; Tulkens et al., 2020). Based on the assumption that most of the microbiome in the semi-naturalistic housing environments at the Emory National Primate Research Center would be dependent on the diet that is controlled across all animals, reassuringly, we found a high degree of convergence in the microbial species represented in RNA contents of EVs extracted from CONT and MALT groups. Also giving us confidence in our results is the predominance of Actinomycetia, Alphaproteobacteria, Betaproteobacteria and Gammaproteobacteria in both groups which agrees with previous profiling of the oral, anal and vaginal microbiome in non-captive rhesus macaques (Chen et al., 2018).

One possible explanation for the high level of alignment of EV-located RNA with sequences of the microbiota may be a skewed representation of EVs that may bleb off from cells in the gastrointestinal tract. To address this possibility, we sought to compare and contrast the RNA found in plasma-derived EVs and 16S rRNA sequencing data obtained from stool samples, a proxy of gut-derived microbiota. While we did note some overlap in the most abundant microbiome-associated phyla between these samples, the proportions were dramatically different. While the gut is the main habitat and source of microbiota, other body parts including cells in the skin, hair, mouth, nose, lung, and genitals (Blum, 2017; Gilbert et al., 2018; Ley et al., 2011; Marsland & Gollwitzer, 2014; Rinninella et al., 2019) have their unique microbiome profile. Therefore, the discrepancy that we see between the proportions of the most abundant phyla in plasma-derived EVs and the stool-derived samples could be reflective of different sources of origins of the microbiome-associated signatures. Our observations would suggest that the microbiome-associated RNA content in circulating EVs are a unique combination of RNA signatures released into the blood stream from the whole body and not just the gut. This interpretation serves to reinforce the concept that RNA in EVs serve as a biological index of biological functions that are occurring throughout the organism.

In thinking of factors that could contribute to the unique microbial species that we found in EVs from both the CONT and MALT groups despite the controlled housing and dietary environments of these animals, transfer through the birth canal, perturbations to epithelial lining of tissues, and differences in available nutrition for the microbiome are worthy of discussion. First, among the earliest determinants of microbiome signatures is exposure to the maternal vaginal microbiome (Dominguez-Bello et al., 2010; Dunn et al., 2017; Ferretti et al., 2018; Jašarević et al., 2015; Mortensen et al., 2021; Mueller et al., 2015). With maltreatment in rhesus macaques propagated through matrilines, an infant that experiences maltreatment is born to a mother that had also experienced maltreatment. Therefore, the unique signatures of microbiome that we observe in CONT and MALT EVs could be a fingerprint of the CONT- and MALT-specific maternal vaginal microbiome that the infant had been exposed to at the time of vaginal birth. Second, another consequence of maltreatment is the development of a leaky gut (Camilleri & Gorman, 2007; Maes et al., 2012; Pohl et al., 2017; Tan et al., 2021). This dimension of digestive biology could result in differential nutrition for bacterial and viral fauna and perturb the balance of the microbiome from “steady state” CONT levels. Additionally, contributing to a leaky gut is a porous epithelium lining the intestinal lumen. Such porosity could extend to epithelia linings of cells and tissues in other parts of the body. From this leakiness of epithelia linings, various combinations of microbiome-associated RNA could be found in circulating EVs. Third, our analyses of community diversity in CONT and MALT samples is striking in that the MALT group has a higher abundance of RNA than the CONT group despite the total number of species being constant between groups. One source of these differences may be that the MALT and CONT animals have a different nutrient microcosm that supports the thriving of different microbial communities and transcription of more RNA. While the exact source of the RNA signatures of unique microbial species in EVs obtained from the CONT and MALT groups cannot definitively be identified, that they exist in the MALT group is suggestive of the influence of maltreatment on the microbiome. Finally, the microbiome of infants and adolescents is responsive to early nutrition by way of being fed breast milk or not (Azad et al., 2013; Gregory et al., 2016; S. Y. Kim & Yi, 2020; Rautava, 2016), and the timing of this feeding from birth (Kim et al., 2019; Pannaraj et al., 2017). However, we posit that breast milk is not a key contributor to the differences in EV-associated microbiome signatures because CONT and MALT animals do not receive different profiles of nursing and have comparable physical growth suggesting no differences in access to nutrition (McCormack et al., 2022).

While an altered microbiome reflected in the RNA profile of circulating plasma EVs in the aftermath of infant maltreatment might seem inconsequential, recent studies of the microbiome and its influence on behavior, physiology and disease counter this perspective. First, the microbiome has been shown to change in response to ELA in rodents and in humans (Cong et al., 2015; Dong & Gupta, 2019; Hantsoo & Zemel, 2021; Kemp et al., 2021; O’Mahony et al., 2009). Second, the microbiome can modulate neurotransmitter abundance and in doing so alter chemical activity in the brain (Berding et al., 2016; Cenit et al., 2017; Chen et al., 2021; Hasan Mohajeri et al., 2018; Jameson & Hsiao, 2018). Third, metabolites that result from microbiome-assisted chemical reactions find their way into circulation and can alter physiology and behavior (Jameson & Hsiao, 2018; Johnson & Foster, 2018; Kim, 2018; Levy et al., 2017; Sampson & Mazmanian, 2015; Sharon et al., 2014; Vuong et al., 2017). Therefore, the differences in microbiome-associated RNA signatures between CONT and MALT samples that we find in circulating plasma EVs could indicate a deeper influence of maltreatment on the microbiome and one more avenue by which maltreatment can exert influence on biological processes in its recipients.

To our knowledge, this is the first time that an EV-focused approach has been used in any species to identify EV-contained signatures of ELA. Released from cells and coursing through the circulatory system, EVs carrying molecular cargo that serve as inter-cellular and even inter-organ messengers of biological information (Braicu et al., 2014; Coakley et al., 2015; Colombo et al., 2014; Isola & Chen, 2017; Raposo & Stoorvogel, 2013; Schorey et al., 2015; Srinivasan et al., 2016; Stahl & Raposo, 2018; Sun et al., 2013). While our findings point us in the direction of potential biomarkers and mediators of infant maltreatment on physiology and behavior, they are accompanied by the caveats of our small sample sizes and that we do not know the cellular origin of these host and microbial RNAs. As this field of inquiry matures, identifying which cells might be most influential in contributing to RNA cargo of EVs in the aftermath of infant maltreatment will shed nuance on the cells, tissues and organ systems that bear the brunt of such ELA. From our approach, we conclude that the effects of ELA can be profiled during adolescence by changes in RNA in EVs that are related to immune function, cellular energetics and microbiome diversity.

## Acknowledgements

The authors want to thank Anne Glenn, Christine Marsteller, Dora Guzman, and the staff at the AAALAC accredited Emory National Primate Research Center (ENPRC) for the excellent technical support and animal care provided during these studies. This study was supported by NIH funding: DA038588, DA052909, MH078105 to MMS, MH128427 to BGD, the Office of Research Infrastructure Programs/OD grant OD11132 (ENPRC Base grant), and an ENPRC Pilot Research Program Grant (ENPRC Base Grant, P51-OD011132) to BGD. BGD is also supported by Department of Pediatrics at USC Keck School of Medicine, the Division of Endocrinology at Children’s Hospital Los Angeles, the Developmental Neuroscience and Neurogenetics Program at The Saban Research Institute, and the Canadian Institute for Advanced Research.

## Supplementary Figs

**Fig. S1:**
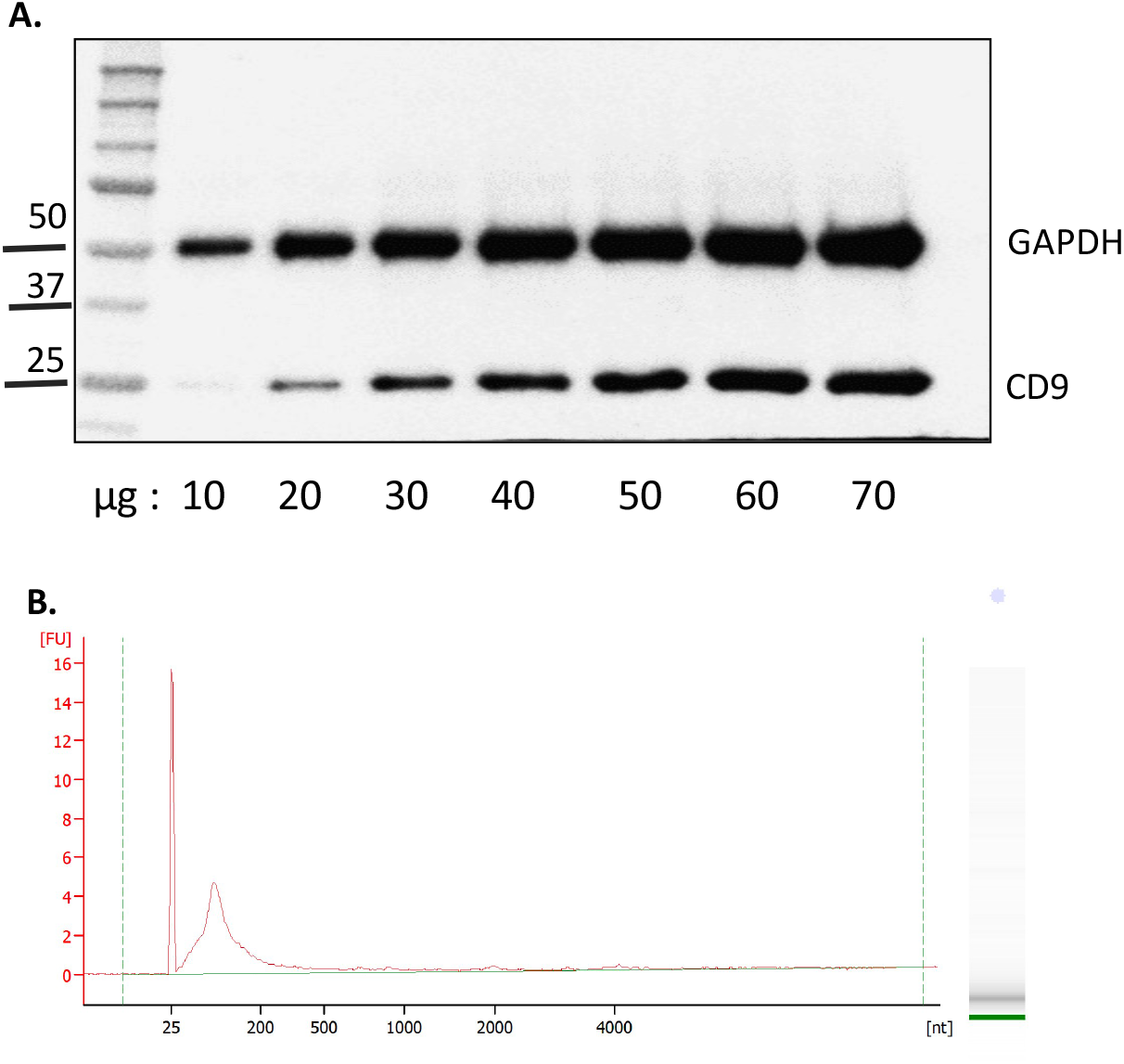
EV isolation and RNA extraction from EVs. **(A)** Western blot showing detection of CD9 biomarker of EVs and GAPDH after protein extraction from isolated EVs. Each lane represents increasing concentration of protein loaded. **(B)** Representative bioanalyzer trace of RNA extracted from EVs.

**Fig. S2:**
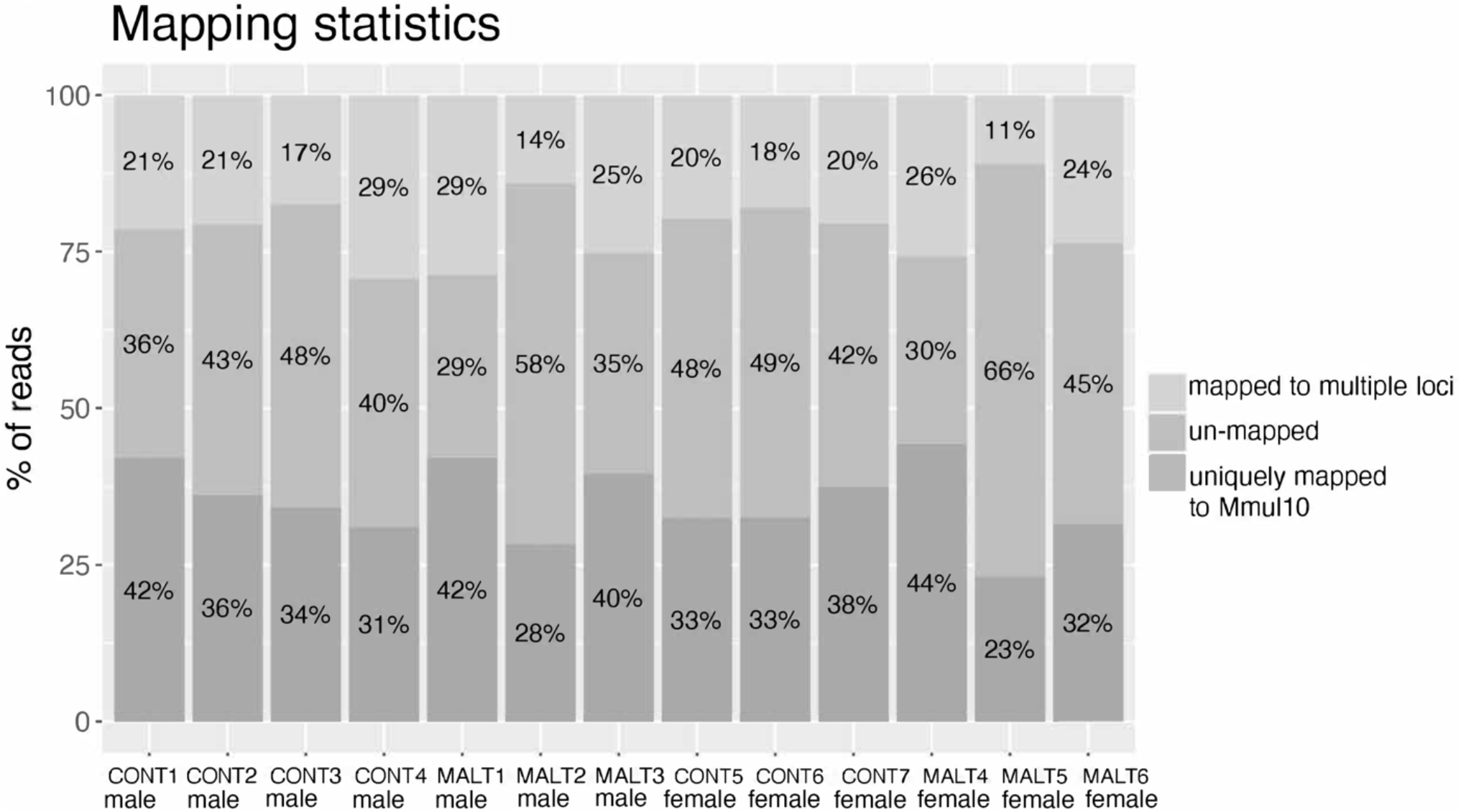
Mapping statistics of EVs RNA-seq. Plot shows the summary of RNA-seq mapping statistics to the reference Mmul10 genome. Three groups of reads that were uniquely mapped, mapped to multiple loci, and not mapped to the reference genome are color-coded. Fractions are given for each sample.

**Fig. S3:**
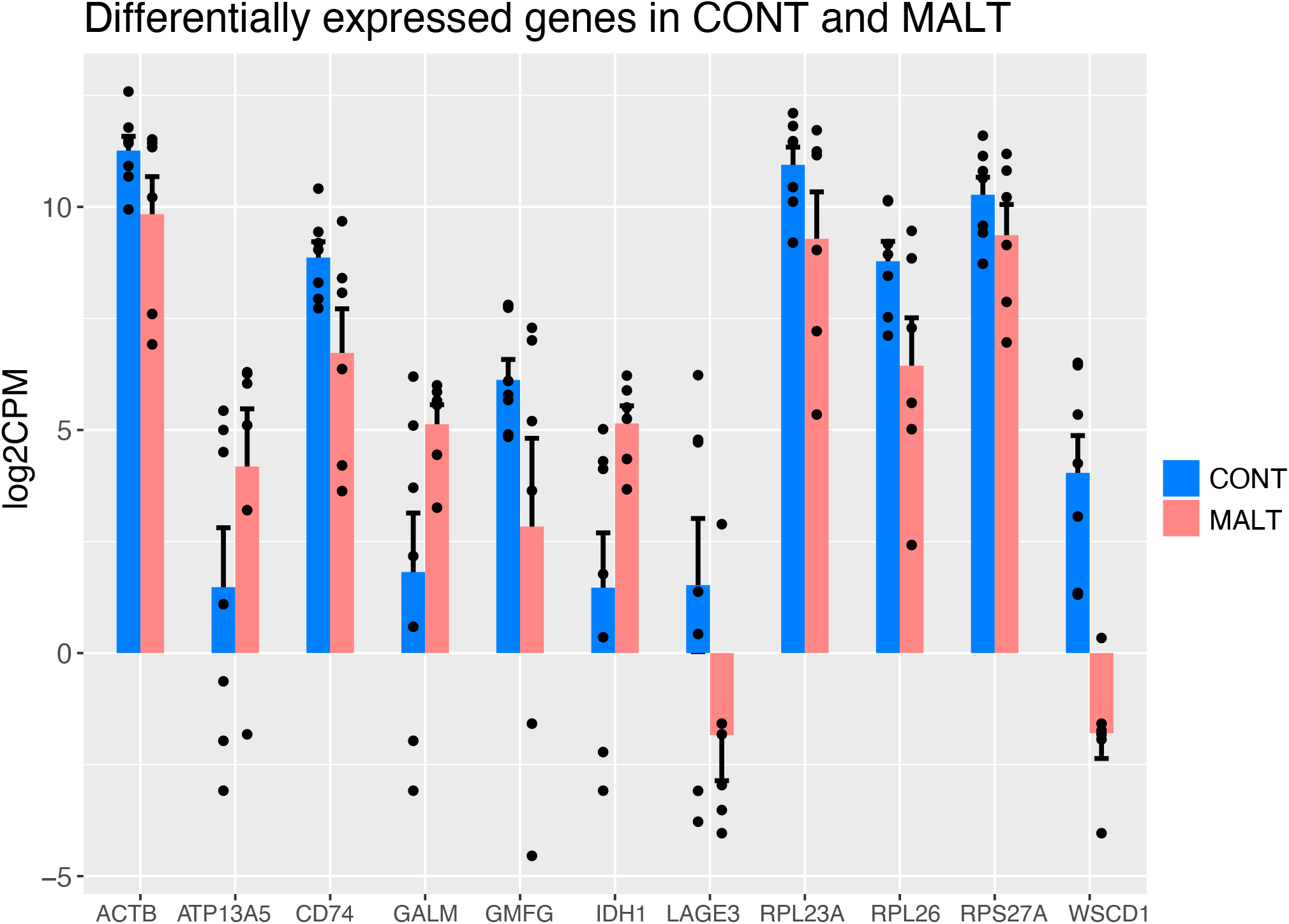
Differentially expressed genes in CONT and MALT. The barplot displays the mean log2CPM (log2 of counts per million) gene expression for each of the eleven genes on x-axis that differentially expressed in CONT and MALT groups (p.adj <0.05). Error bars show the standard error of the mean. The scatter plot shows the individual data points in black. The colors of the bars are coded to represent two groups of comparison. The log2CPM values were obtained from voom function of limma package in R.

**Fig. S4:**
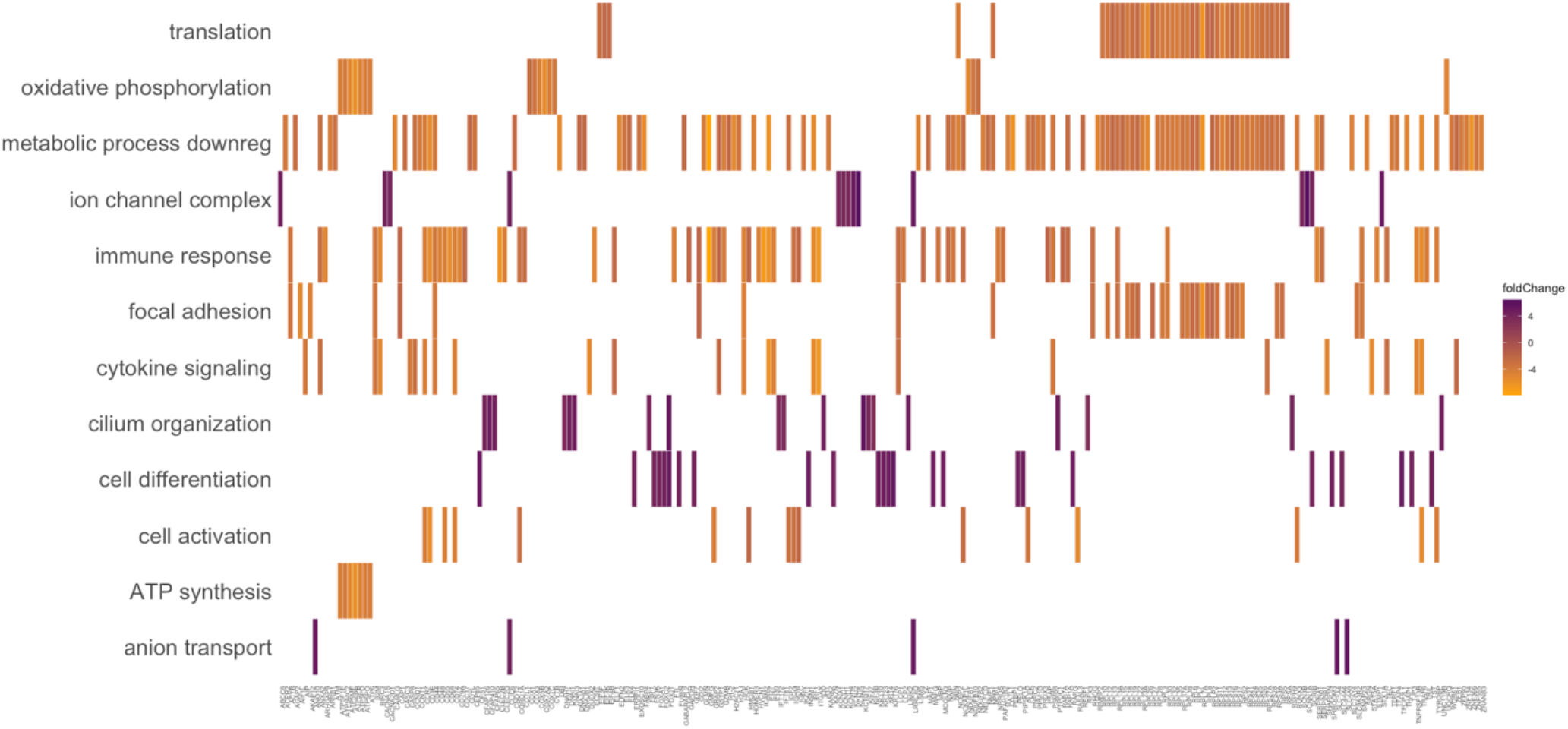
Perturbations of genes pathways involved in specific biological processes by infant maltreatment are evident from RNA signatures found in circulating EVs. Plot showing the composition of significantly enriched Gene Ontology terms from BP (biological process) category. Name of the terms are plotted on the y-axis. List of the genes is plotted on the x-axis. Log2 fold change is color-coded with gradient from brown (downregulated in MALT) to purple (upregulated in MALT).

**Fig. S5:**
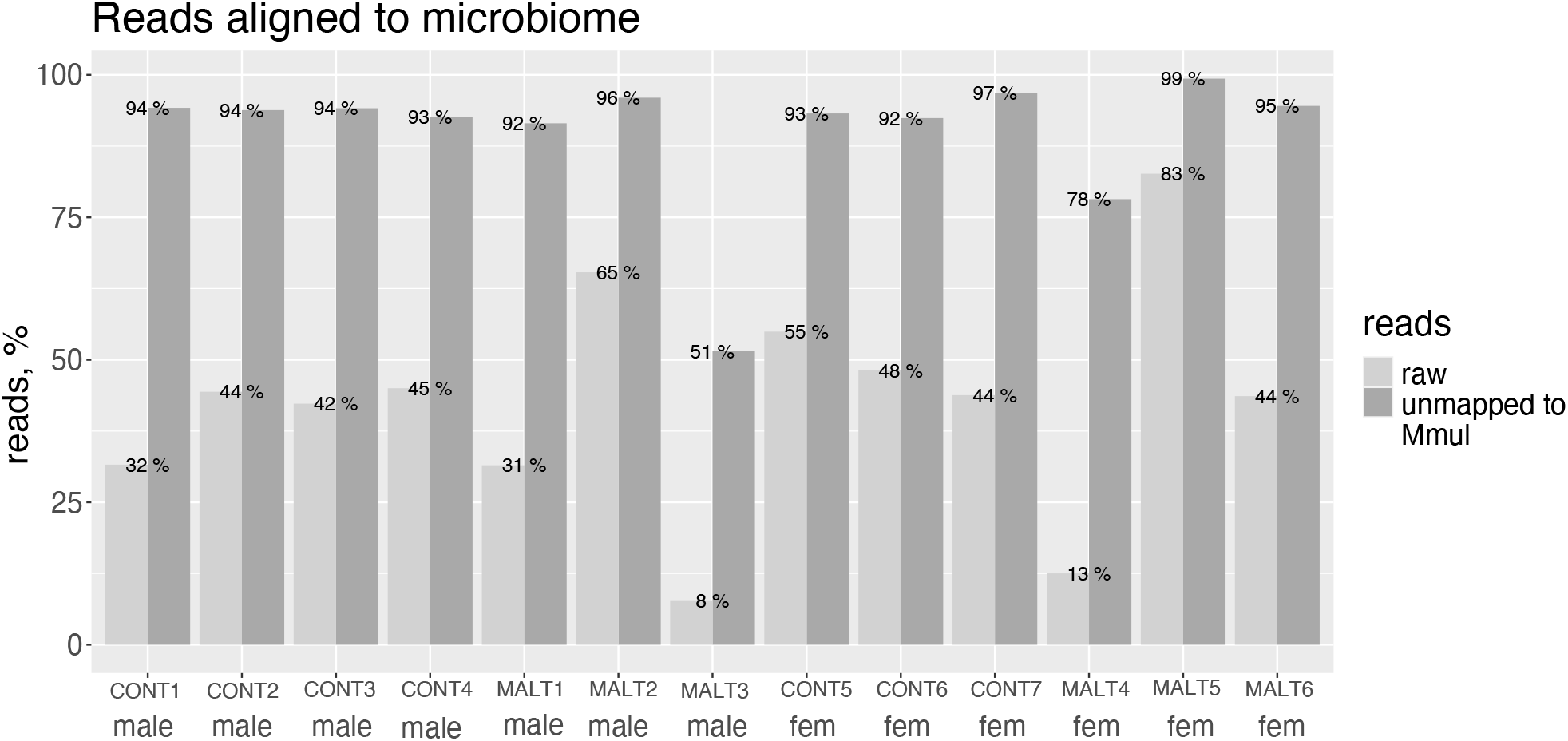
RNA in EVs that do not align to the macaque reference genome align to the microbiome meta-transcriptome. Reads that did not align to the reference Mul10 genome were identified as bacterial and viral with the Kraken algorithm. The plot shows the percentage of reads belonging to microbiome in each sample before and after filtering. Light grey colored bars show the percentage of raw reads that we aligned to microbiome reference genome; dark grey colored bars represent percentage of reads that were not aligned to reference Macaque genome, and are detected as microbiome-derived RNA reads.

**Fig. S6:**
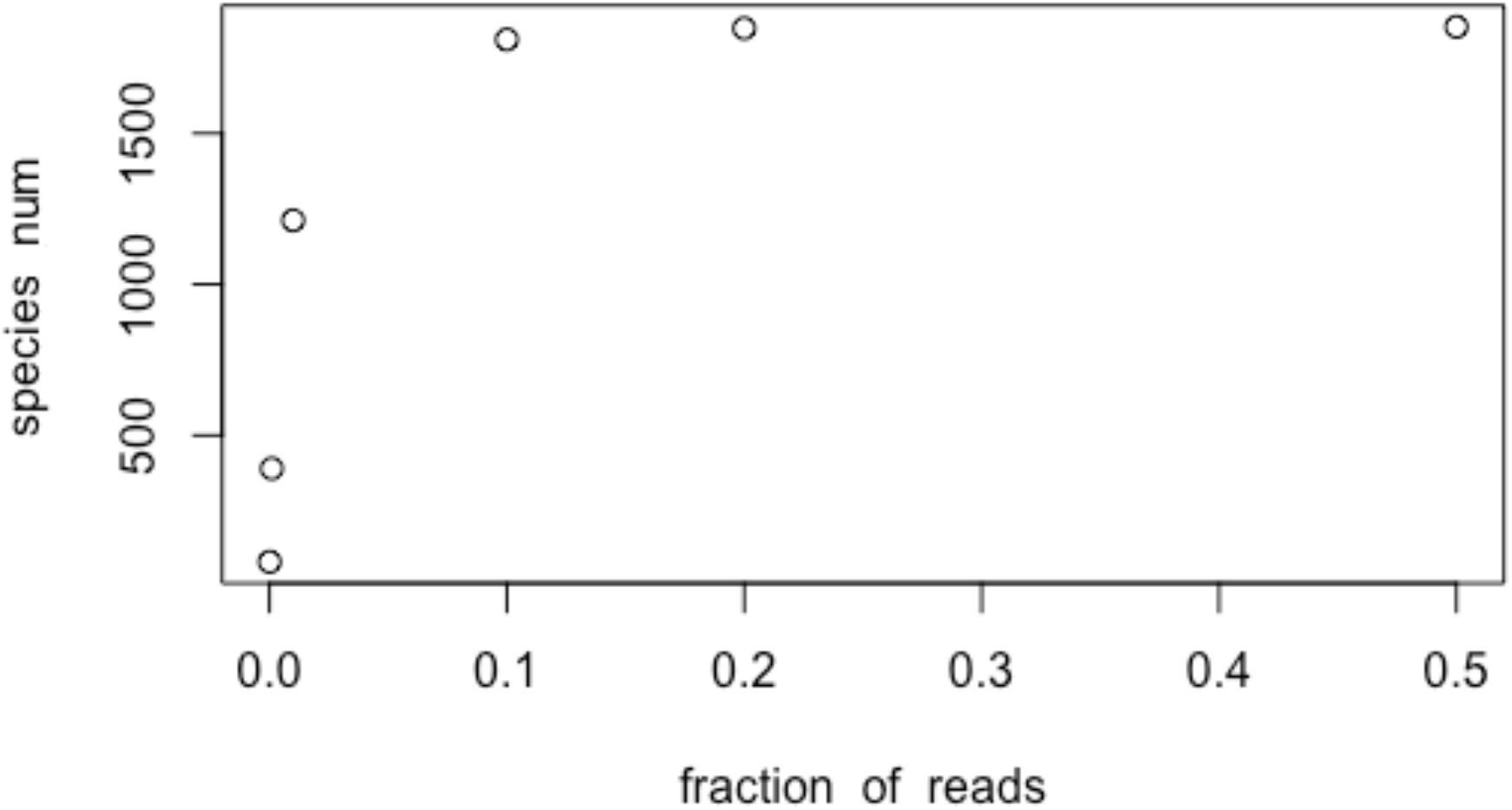
RNA in EVs aligning to the microbiome meta-transcriptome represent a significant portion of the microbiome. Plot illustrating the quality and the stability of the data. The rarefaction curve revealed that the rarest species would remain to be sampled with only 20% of reads. Even 10% of randomly taken reads detects most species found with all available reads.

**Fig. S7.**
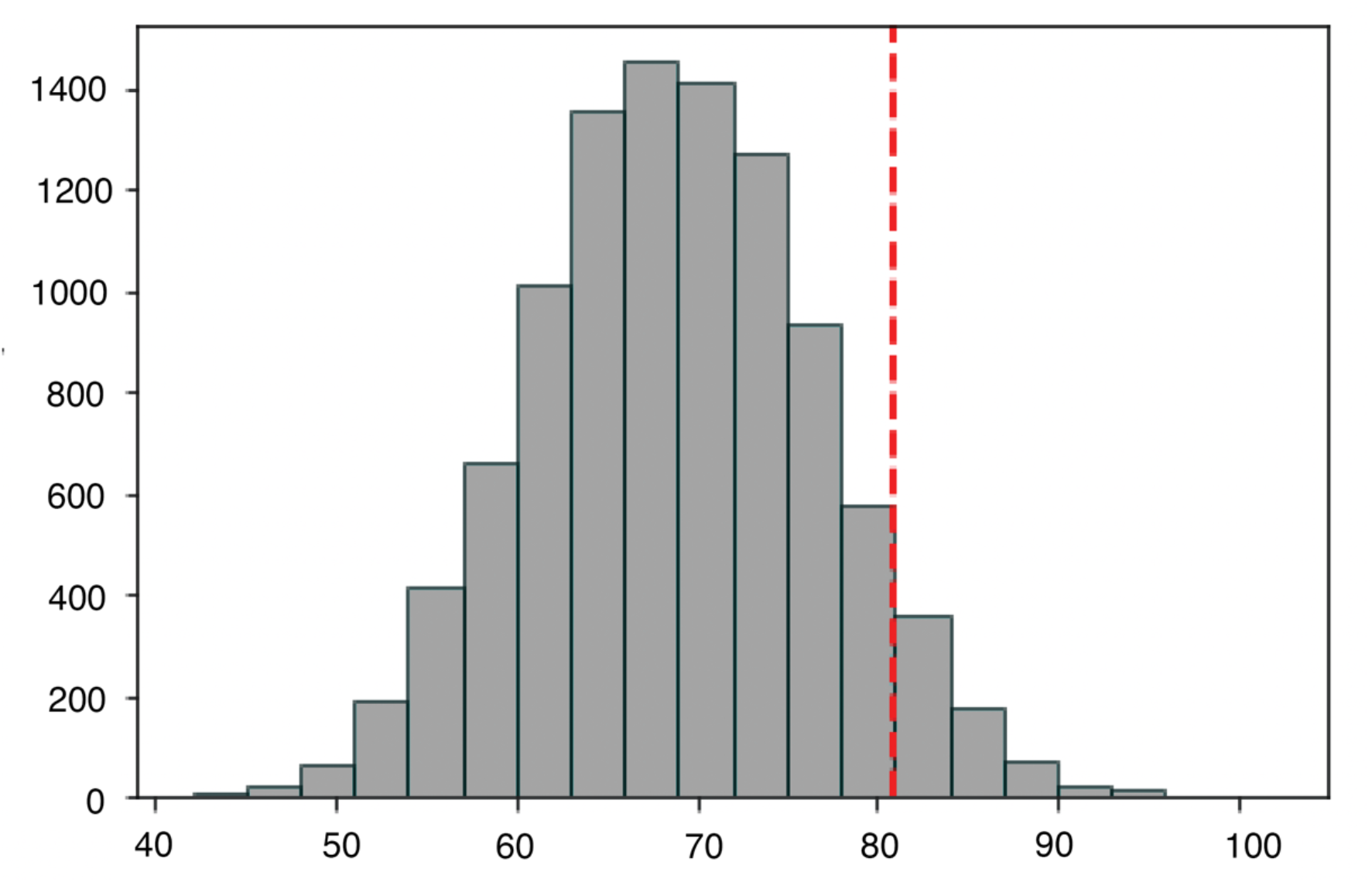
Permutation test. Visual representation of the distribution of 5000 values from the permutation test, measuring the expected difference in species count between MALT and CONT groups. For each species majority was determined by absolute (n of presence in CONT - n of presence in MALT) > 4. The observed measure (red line) of species count difference falls outside the range of expected values, yielding a p-value of 0.0478 indicating strong evidence against the null hypothesis of no difference.

**Fig. S8:**
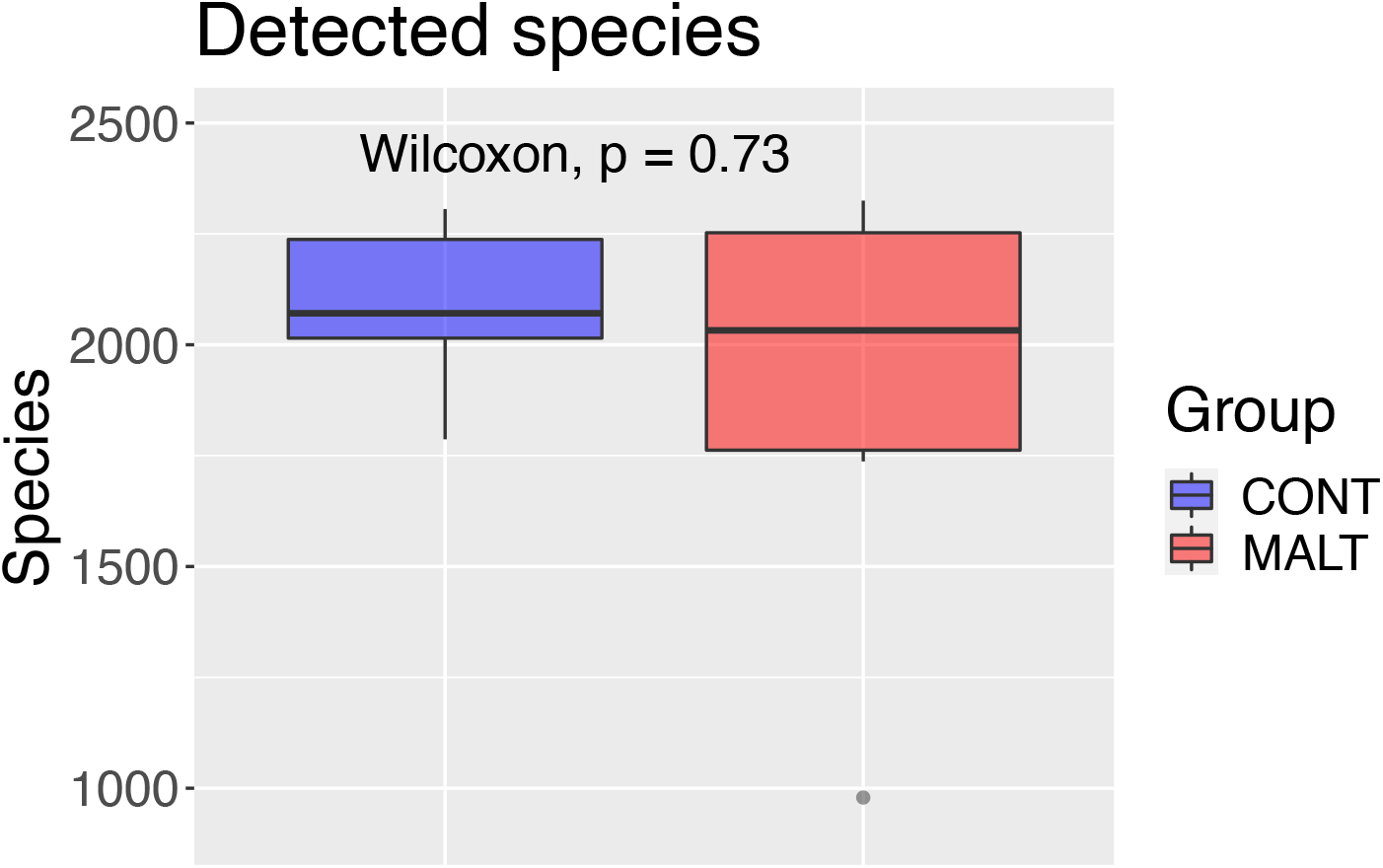
Infant maltreatment does not significantly change the total number of microbiome species as reflected in RNA content of EVs. Plot showing the total number of microbiome species in MALT (red) and CONT (blue) groups as reflected by alignment of RNA in EVs to the microbiome meta-transcriptome. Boxplot represents the median (line in the middle of the box), upper 25% quantile, lower 25% quantile, and the interquartile ranges (upper and lower whiskers). Wilcoxon test derived p-value is shown.

**Table S1:** List of GO terms in the functional category “Biological processes” that are differentially enriched in plasma-derived EVs of MALT animals with their adjusted p-values.

| **ID** | **Description** | **p.adjust** |
| --- | --- | --- |
| GO:0006413 | translational initiation | 2.42E-08 |
| GO:0006613 | cotranslational protein targeting to membrane | 2.42E-08 |
| GO:0006614 | SRP-dependent cotranslational protein targeting to membrane | 2.42E-08 |
| GO:0019080 | viral gene expression | 2.42E-08 |
| GO:0019083 | viral transcription | 2.42E-08 |
| GO:0045047 | protein targeting to ER | 2.42E-08 |
| GO:0070972 | protein localization to endoplasmic reticulum | 2.42E-08 |
| GO:0072599 | establishment of protein localization to endoplasmic reticulum | 2.42E-08 |
| GO:0000184 | nuclear-transcribed mRNA catabolic process, nonsense-mediated decay | 4.13E-08 |
| GO:0006612 | protein targeting to membrane | 5.99E-07 |
| GO:0044419 | biological process involved in interspecies interaction between organisms | 1.24E-06 |
| GO:0000956 | nuclear-transcribed mRNA catabolic process | 2.42E-06 |
| GO:0090150 | establishment of protein localization to membrane | 0.00046 |
| GO:0072594 | establishment of protein localization to organelle | 0.00063 |
| GO:0002181 | cytoplasmic translation | 0.00063 |
| GO:0006605 | protein targeting | 0.00083 |
| GO:0006412 | translation | 0.00108 |
| GO:0022613 | ribonucleoprotein complex biogenesis | 0.00232 |
| GO:0006402 | mRNA catabolic process | 0.0027 |
| GO:0006401 | RNA catabolic process | 0.0030 |
| GO:0043604 | amide biosynthetic process | 0.00300 |
| GO:0006952 | defense response | 0.00332 |
| GO:0043043 | peptide biosynthetic process | 0.00357 |
| GO:0050867 | positive regulation of cell activation | 0.00359 |
| GO:0019221 | cytokine-mediated signaling pathway | 0.00465 |
| GO:0016032 | viral process | 0.00501 |
| GO:0034655 | nucleobase-containing compound catabolic process | 0.00501 |
| GO:0044403 | biological process involved in symbiotic interaction | 0.00545 |
| GO:0016071 | mRNA metabolic process | 0.00754 |
| GO:0019439 | aromatic compound catabolic process | 0.00778 |
| GO:0042254 | ribosome biogenesis | 0.00778 |
| GO:0009607 | response to biotic stimulus | 0.00778 |
| GO:0043207 | response to external biotic stimulus | 0.00778 |
| GO:0051707 | response to other organism | 0.00778 |
| GO:0002696 | positive regulation of leukocyte activation | 0.00780 |
| GO:0051251 | positive regulation of lymphocyte activation | 0.00780 |
| GO:0010605 | negative regulation of macromolecule metabolic process | 0.00780 |
| GO:1901361 | organic cyclic compound catabolic process | 0.00780 |
| GO:0071345 | cellular response to cytokine stimulus | 0.01027 |
| GO:0072657 | protein localization to membrane | 0.01192 |
| GO:0042273 | ribosomal large subunit biogenesis | 0.01601 |
| GO:0019752 | carboxylic acid metabolic process | 0.01771 |
| GO:0001775 | cell activation | 0.01771 |
| GO:0009892 | negative regulation of metabolic process | 0.01796 |
| GO:0043436 | oxoacid metabolic process | 0.01821 |
| GO:0050870 | positive regulation of T cell activation | 0.01835 |
| GO:0007389 | pattern specification process | 0.01855 |
| GO:0098542 | defense response to other organism | 0.01860 |
| GO:0006082 | organic acid metabolic process | 0.01860 |
| GO:0046700 | heterocycle catabolic process | 0.01860 |
| GO:0007007 | inner mitochondrial membrane organization | 0.01860 |
| GO:0015985 | energy coupled proton transport, down electrochemical gradient | 0.01860 |
| GO:0015986 | ATP synthesis coupled proton transport | 0.01860 |
| GO:0042407 | cristae formation | 0.01860 |
| GO:0042776 | mitochondrial ATP synthesis coupled proton transport | 0.01860 |
| GO:0006955 | immune response | 0.01958 |
| GO:0046649 | lymphocyte activation | 0.02102 |
| GO:0043123 | positive regulation of I-kappaB kinase/NF-kappaB signaling | 0.02130 |
| GO:0030855 | epithelial cell differentiation | 0.02130 |
| GO:0033365 | protein localization to organelle | 0.02130 |
| GO:0050865 | regulation of cell activation | 0.02130 |
| GO:0044270 | cellular nitrogen compound catabolic process | 0.02143 |
| GO:0045321 | leukocyte activation | 0.02241 |
| GO:0002682 | regulation of immune system process | 0.02494 |
| GO:0032943 | mononuclear cell proliferation | 0.03149 |
| GO:0046651 | lymphocyte proliferation | 0.03149 |
| GO:0002684 | positive regulation of immune system process | 0.03327 |
| GO:0044782 | cilium organization | 0.03799 |
| GO:0015711 | organic anion transport | 0.03807 |
| GO:0006364 | rRNA processing | 0.03852 |
| GO:0006119 | oxidative phosphorylation | 0.03987 |
| GO:0044265 | cellular macromolecule catabolic process | 0.03987 |
| GO:0002252 | immune effector process | 0.04263 |
| GO:0071356 | cellular response to tumor necrosis factor | 0.04263 |
| GO:0002694 | regulation of leukocyte activation | 0.04263 |
| GO:0051249 | regulation of lymphocyte activation | 0.04263 |
| GO:0051649 | establishment of localization in cell | 0.04539 |
| GO:0015698 | inorganic anion transport | 0.04859 |
| GO:0006754 | ATP biosynthetic process | 0.04859 |
| GO:0060271 | cilium assembly | 0.04859 |
| GO:0002376 | immune system process | 0.04859 |
| GO:0032944 | regulation of mononuclear cell proliferation | 0.04859 |
| GO:0050670 | regulation of lymphocyte proliferation | 0.04859 |
| GO:0032946 | positive regulation of mononuclear cell proliferation | 0.04859 |
| GO:0050671 | positive regulation of lymphocyte proliferation | 0.04859 |
| GO:0070665 | positive regulation of leukocyte proliferation | 0.04859 |
| GO:0034097 | response to cytokine | 0.04859 |

**Table S2:**
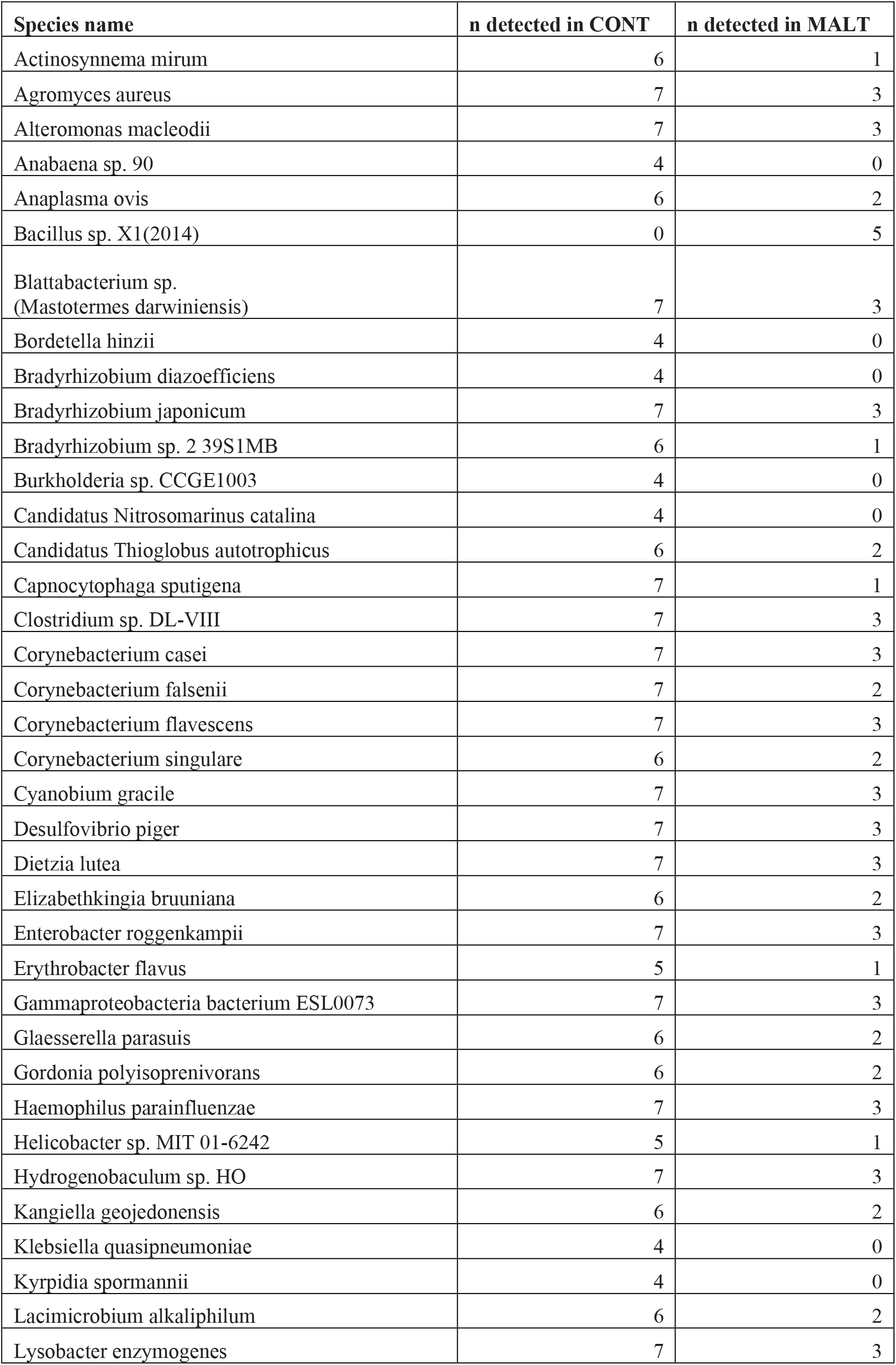
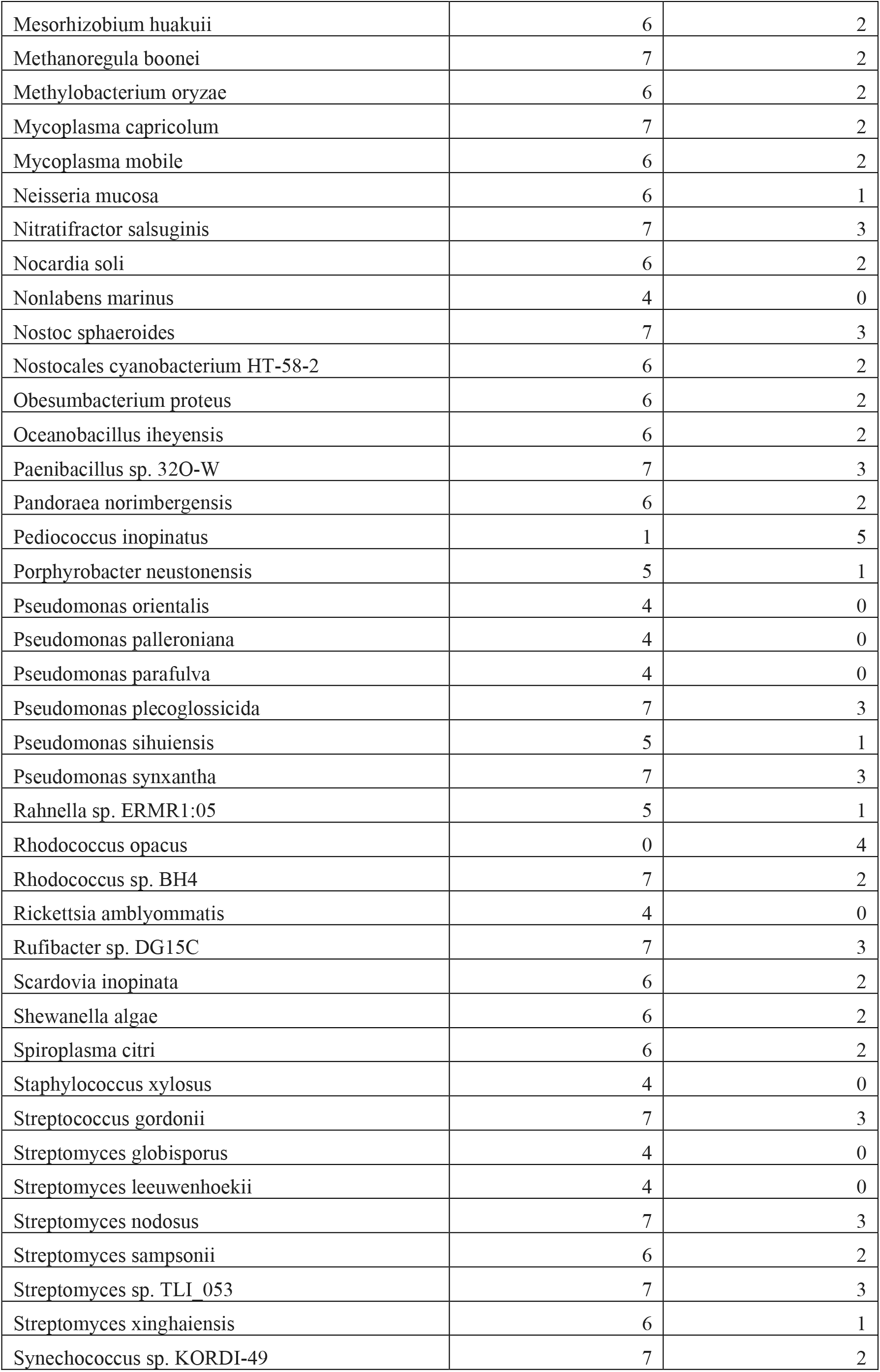
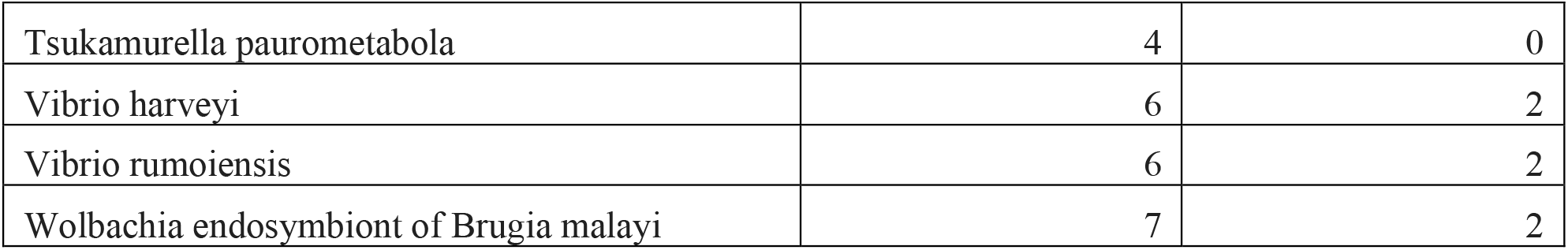
List of microbiome species that were identified in majority of CONT and MALT plasma-derived EVs (present in more than 4 animals in one group but not another).

